# Oscillatory dynamics of perceptual to conceptual transformations in the ventral visual pathway

**DOI:** 10.1101/259127

**Authors:** Alex Clarke, Barry J. Devereux, Lorraine K. Tyler

## Abstract

Object recognition requires dynamic transformations of low-level visual inputs to complex semantic representations. While this process depends on the ventral visual pathway (VVP), we lack an incremental account from low-level inputs to semantic representations, and the mechanistic details of these dynamics. Here we combine computational models of vision with semantics, and test the output of the incremental model against patterns of neural oscillations recorded with MEG in humans. Representational Similarity Analysis showed visual information was represented in alpha activity throughout the VVP, and semantic information was represented in theta activity. Furthermore, informational connectivity showed visual information travels through feedforward connections, while visual information is transformed into semantic representations through feedforward and feedback activity, centered on the anterior temporal lobe. Our research highlights that the complex transformations between visual and semantic information is driven by feedforward and recurrent dynamics resulting in object-specific semantics.

## Introduction

Visual object recognition requires dynamic transformations of information from low-level visual inputs to higher-level visual properties and ultimately complex semantic representations. These processes rely on the ventral visual pathway (VVP) from the occipital lobe along the ventral surface of the temporal lobe (Kravitz et al., 2013), with the perirhinal cortex (PRC) sitting at the apex of the pathway (Barense et al., 2012; Bussey and Saksida, 2002; Clarke and Tyler, 2014; Cowell et al., 2010; Taylor et al., 2006; Tyler et al., 2013). Along the VVP, object representations become increasingly complex and abstracted from their inputs, such that higher-level visual properties are coded in LOC and posterior IT that show object invariance (DiCarlo et al., 2012; Kravitz et al., 2013), alongside conceptual properties of objects that are sufficient to distinguish between different superordinate categories (Tyler et al., 2013). In contrast, object-specific semantic representations are seen in the perirhinal cortex, at the most anterior part of the VVP, which is hypothesised to form complex conjunctions of properties from more posterior regions to enable fine-grained distinctions between conceptually similar objects (Barense et al., 2012; Bussey and Saksida, 2002; Clarke and Tyler, 2014; Cowell et al., 2010; Kivisaari et al., 2012; Tyler et al., 2004, 2013). Yet this view of recognition - where both activity and the complexity of object information progresses along the posterior to anterior axis in the VVP - is fundamentally incomplete as it does not take into account the temporal dynamics of feedforward and feedback processes and their interactions.

The brain’s anatomical structure suggests that complex interactions between bottom-up and top-down processes must be a key part of object processing, as demonstrated by the abundance of lateral and feedback anatomical connections within the VVP and beyond (Bullier, 2001; Lamme and Roelfsema, 2000). Research using time-resolved imaging methods have shown that both feedforward and recurrent dynamics in the VVP underpin object representations, where visual inputs activate semantic information within the first 150 ms, and object-specific semantic representations emerge beyond 200 ms supported by recurrent activity between the ATL and posterior VVP (Chan et al., 2011; Clarke et al., 2011, 2013, 2015; Poch et al., 2015; Schendan and Maher, 2009).

While this research provides spatial and temporal signatures of the fundamental aspects of recognition – namely visual and semantic processing - two important limitations remain. First, research tends to focus on three aspects of objects – low-level visual properties, superordinate category information (e.g. animals, tools, animate/inanimate) and object-specific semantics (e.g. tiger, hammer). This paints a compartmentalised picture that fails to capture the incremental transitions whereby vision seamlessly activates meaning. Second, while there is increasing knowledge of the oscillatory mechanisms underpinning basic visual processing (Jensen et al., 2014; Tallon-Baudry and Bertrand, 1999), models of how visual inputs activate meaning lack such detail. Here, we overcome these limitations by combining current computational models of vision with a model of semantics, to obtain quantifiable estimates of the incremental representations from low-level visual inputs to complex semantic representations. This model is then tested against neural activity using Representational Similarity Analysis (RSA) (Kriegeskorte et al., 2008; Nili et al., 2014) to reveal how oscillatory activity along the VVP codes for visual and semantic object properties.

Deep neural networks (DNNs) have proved highly successful for vision, both to provide an engineering solution to labelling objects (Krizhevsky et al., 2012) and to map the outputs from the DNN to brain representations of objects in space and time (Cichy et al., 2014, 2016; Devereux et al., under review; Güçlü and Gerven, 2015; Seeliger et al., 2017). DNNs for vision are composed of multiple layers, and as the layers progress, the nodes become sensitive to more complex, higher-level visual image features in a similar progression to the human ventral visual pathway (Cichy et al., 2014, 2016; Devereux et al., under review; Güçlü and Gerven, 2015). However, current DNNs tell us little about an object’s *semantic* representation. While visual DNNs can provide accurate labels for images, the output layers do not capture how different objects are related in meaning. This is revealed by recent fMRI research, showing that while DNNs explain visual processes in the posterior vemtral temporal cortex (pVTC), additional semantic computational models are required to capture the semantic information about object representations in the pVTC and PRC (Devereux et al., under review). This work used a recurrent attractor network (AN) for object semantics, as ANs have been shown to capture how objects semantically relate to one another (Cree et al., 1999, 2006; Devereux et al., 2015). This occurs because the activation across the nodes in the model capture the activation of different semantic features (such as ‘is round’, ‘has a handle’, ‘is thrown’, etc). Further, the dynamics of how these nodes become activated mirrors both behavioural responses and MEG time-courses during object recognition (Clarke et al., 2013; Devereux et al., 2015; Randall et al., 2004). Together, the DNN and AN provide complementary aspects of object recognition. As in Devereux et al (under review), by using the output of the visual DNN as input into the semantic AN, we further provide a potential route by which visual representations can directly activate semantic knowledge. Most importantly, however, combining the DNN and AN gives us a quantifiable computational approach that models the incremental visual and semantic properties of objects, from low-level vision to high level semantics. This approach can be combined with RSA for dynamic measures of brain activity to show how different types of visual and semantic information are coded in dynamic patterns of brain activity along the VVP.

The brain activity we focus on here are neural oscillations. Oscillations are a ubiquitous property of the brain, and are known to be modulated by various aspects of vision and memory in humans (Fell and Axmacher, 2011; Hanslmayr et al., 2012; Helfrich and Knight, 2016; Jensen et al., 2014; Watrous et al., 2015a). Recent studies have begun to show how the ongoing phase of an oscillation can be used to decode specific stimuli (Lopour et al., 2013; Ng et al., 2013; Schyns et al., 2011; Turesson et al., 2012; Watrous et al., 2015b). These studies have shown that frequency-specific activity can be used to decode the specific features of visual objects, or object categories, suggesting that oscillatory phase could provide a mechanism for encoding stimulus information within a region. Further, oscillations may help to coordinate the activity between regions in the VVP enabling object information to be transformed over space and time as meaning is accessed from vision.

Here, we combine RSA with neural oscillations and computational models, that could provide an important advance in determining the dynamic flow of different types of object information during recognition – both in terms of how different regions represent visual and semantic information and how information is transformed across regions. To achieve this, we recorded MEG while participants viewed a large set of common objects from diverse superordinate categories. The combined DNN and AN models provided predictions for how objects should be similar to one another, and these predictions were tested against the MEG data using RSA (Figure 1). The MEG signals were source localized, and single object oscillatory phase patterns were extracted from 5 regions of interest in the VVP. Based on these phase patterns across objects, we could determine how similar objects were to each other within each ROI, and track this over time and frequency. RSA then allows us to test the degree to which the object similarity according to the computational model is reflected in oscillatory phase signals over space, time and frequency. We predict both a spatial and temporal hierarchy in the VVP between visual object information and semantics. Crucially, recurrent activity will be associated with the activation of semantic object information that will also depend on the coordinated activity within the VVP. Whilst visual object properties are predicted in alpha (VanRullen et al., 2014), semantic information may be more associated with theta activity (Halgren et al., 2015) and gamma activity (Mollo et al., 2017; Supp et al., 2007).

**Figure 1.**
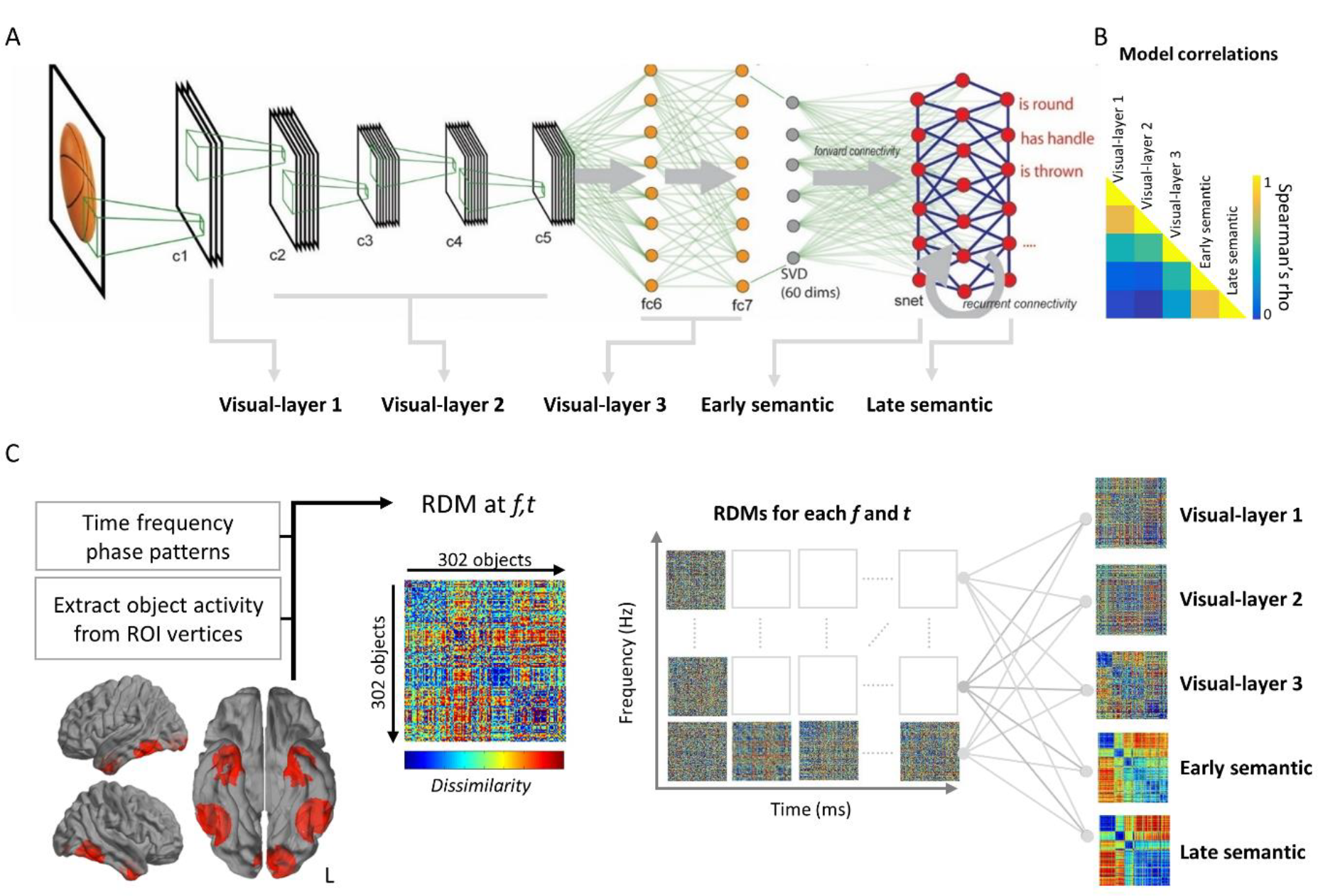
Representational similarity analysis (RSA) using computational models and oscillations. (A). The combined visual DNN and semantic AN models the low-level visual properties of the input, and higher-level image properties that increase in complexity across layers c1 to fc7. The visual properties in fc7 then map onto a recurrent AN that activates the semantic features associated with the input. In our analyses, we combined layers of the DNN into three visual model RDMs, and combined the AN into two semantic model RDMs capturing increasingly specific visual and semantic information. (B). Correlations between the different visual and semantic model RDMs. (C). RSA analysis of time-frequency data. Spatio-temporal activity patterns are extracted from an ROI for each object. Time-frequency phase is calculated for each ROI, and RDMs are created for each point in time and for each frequency. Each RDM is then correlated with each RDM from the computational model to test when and at what frequency different object properties are represented in oscillatory phase patterns. The procedure is then repeated for all ROIs.

## Methods

We re-analysed MEG data reported in Clarke et al., (2015), and thus only summarise the main aspects of study design here.

### Participants & procedure

Fourteen participants took part in the study. Two participants were excluded from the analysis due to poor source reconstruction results (failure to show occipital activity ~100 ms after onset of object) leaving twelve participants in the analysis. Participants performed a basic-level naming task (e.g. ‘tiger’) with 302 common objects from a diverse range of superordinate categories including animals, clothing, food, musical instruments, tools and vehicles. All objects were presented in color as single objects on a white background. Each trial began with a black fixation cross on a white background for 500 ms before the object was shown for 500 ms, and followed by a blank screen lasting between 2400 and 2700 ms. The order of stimuli was pseudo-randomized.

### MEG/MRI recording

Continuous MEG data were recorded using a whole-head 306 channel (102 magnetometers, 204 planar gradiometers) Vector-view system (Elekta Neuromag, Helsinki, Finland) located at the MRC Cognition and Brain Sciences Unit, Cambridge, UK. Eye movements and blinks were monitored with electro-oculogram (EOG) electrodes placed around the eyes, and five Head-Position Indicator (HPI) coils were used to record the head position (every 200 ms) within the MEG helmet. The participants’ head shape was digitally recorded using a 3D digitizer (Fastrak Polhemus Inc., Colchester, VA), along with the positions of the EOG electrodes, HPI coils, and fiducial points. MEG signals were recorded at a sampling rate of 1000 Hz, with a band-pass filter from 0.03 to 125 Hz. To facilitate source reconstruction, 1 mm3 T1-weighted MPRAGE scans were acquired during a separate session with a Siemens 3T Tim Trio scanner (Siemens Medical Solutions, Camberley, UK) located at the MRC Cognition and Brain Sciences Unit, Cambridge, UK.

### MEG preprocessing

Initial processing of the raw data used MaxFilter version 2.0 (Elektra-Neuromag) to detect bad channels that were subsequently reconstructed by interpolating neighboring channels. The temporal extension of the signal-space separation technique was applied to the data every 10 seconds in order to segregate the signals originating from within the participants’ head from those generated by external sources of noise. A correlation limit of 0.6 was used as this has been shown to additionally remove noise from close to the head, as produced during speech (Medvedovsky et al., 2009), and head movement compensation was applied. The resulting MEG data were low-pass filtered at 200 Hz in forward and reverse directions using a 5th order Butterworth digital filter, high-pass filtered at 0.1 Hz using a 4th order butterworth filter and residual line noise was removed with a 5th order butterworth stop-band filter between 48 and 52 Hz. Data were epoched from −1.5 to 2 seconds, and downsampled to 500 Hz using SPM12 (Wellcome Institute of Imaging Neuroscience, London, UK).

Independent components analysis (ICA) was used to remove artefactual signals, using runica implemented in EEGLab (Delorme and Makeig, 2004) and SASICA (Chaumon et al., 2015). Components of the data that showed a Pearson’s correlation greater 0.4 with either EOG channel were removed from the data, as were components correlated with the ECG recording. SASICA and FASTER were additionally used to identify components related to muscle and high-frequency artefacts, and components that showed a rising profile of evoked activity between 200 ms and 1 second were removed (these characterise speech artefacts). All components were visually inspected to confirm removal, as recommended (Chaumon et al., 2015). ICA was applied to the magnetometers and gradiometers separately. After ICA, a baseline correction was applied to all trials using data from −500 to 0 ms. Items that were incorrectly named were excluded, where an incorrect name was defined as a response that did not match the correct concept.

### Source localisation

Source localisation of MEG signals used a minimum-norm procedure applied in SPM12. First, the participants MRI images were segmented and spatially normalized to an MNI template brain. A template cortical mesh with 8,196 vertices was inverse normalized to the individual’s specific MRI space. MEG sensor locations were coregistered to MRI space using the fiducial points and digitized head-points obtained during acquisition. The forward model was created using the single shell option to calculate the lead-fields for the sources oriented normal to the cortical surface. The data from both magnetometers and gradiometers were inverted together using the group inversion approach to estimate activity at each cortical vertex using a minimum norm solution (IID). A frequency window of 0 to 150 Hz was specified and no hanning window was applied.

### Representational Similarity Analysis (RSA)

RSA was used to compare the similarity/distances between objects based on computational models, and the similarity derived from oscillatory patterns. This requires we calculate representational dissimilarity matrices (RDMs) from both the computational model layers, and from source localised MEG signals.

#### RDMs from computational models

The computational models used here are those that have been successfully used to describe the gradient of visual to semantic object representations along the VVP in fMRI (Devereux et al., under review).

#### Visual Deep Neural Network

We used the deep neural network (DNN) model of Krievhesky et al. (2012), as implemented in the Caffe deep learning framework (Jia et al., 2014), and trained on the ILSVRC12 classification dataset from ImageNet. We used the first 7 layers of the DNN, consisting of five convolutional layers (conv1-conv5) followed by two fully-connected layers (fc6 & fc7). The convolutional kernels learned in each convolutional layer correspond to filters receptive to particular kinds of visual input. In the first convolutional layer, the filters reflect low-level properties of stimuli, and include one sensitive to edges of particular spatial frequency and orientation, as well as filters selective for particular colour patches and colour gradients (Krizhevsky et al., 2012; Zeiler and Fergus, 2014). Later DNN layers are sensitive to more complex visual information, such as the presence of specific visual objects or object parts(e.g. faces of dogs, legs of dogs, eyes of birds & reptiles; see Zeiler and Fergus, 2014), irrespective of spatial scale, angle of view etc. We presented 627 images to the pre-trained network (including the 302 images presented to participants), where each image represented a concept listed in a large property norm corpus (Devereux et al., 2014). This produced activation values for all nodes in each layer of the network for each image.

To create RDMs for each layer of the DNN, we first applied PCA to reduce the dimensionality of each layer while keeping the components that explained 100% of the variance. For example, fc6 has 4096 nodes, and after PCA each of the 627 object images was represented by a 626 length vector. This was found to dramatically improve the relationship between MEG signals and the DNN, which may be because the white space surrounding images was reduced from being represented across a large number of nodes to a few components, meaning the similarity between objects was focussed on the areas of the images where the objects appeared. While the PCA improved the relationship between MEG signals and the DNN for objects isolated from backgrounds, this would not be expected for naturalistic images. Following PCA, we excluded all object activations that were not in the present study, leaving 302.

As many of the layers were highly correlated, and to reduce the number of RDMs tested, subsets of the 7 layers were combined. The object activation matrices were concatenated across layers, and the dissimilarity between network activity for different object images was calculated as 1 - Pearson’s correlation. This was applied to conv1, concatenated activations from conv2-5, and concatenated activations from fc6 and 7, which are referred to as visual-layer 1, 2 and 3 respectively. A model RDM was also created based on concatenated data from all layers of the DNN.

#### Semantic Attractor Network

DNNs have proven effective in labelling object images in complex contexts. However visual DNNs do not capture object semantics, because although they can find the correct labels for images, they do not capture how different objects are semantically related to one another (e.g. that a dog and cat are related in meaning), and only takes into account the similarity of their visual properties, rather than also taking into account non-visual and functional information (Devereux et al., under review). To provide one potential route for the relationship between higher-level visual properties and semantic properties, we use the output from the DNN as input to an attractor network (AN) model of semantics.

Our semantic knowledge of concrete concepts can be captured by distributed semantic feature models (Cree and McRae, 2003; Rogers and McClelland, 2004; Taylor et al., 2011; Tyler and Moss, 2001), where each concept is represented by a set of features – e.g. *is shiny, has a handle, used for chopping* are features of a knife from the property norming corpus of Devereux et al., (2014). Based on semantic features, the similarity between concepts is accounted for on the basis of the features they share, while distinctive features allow for differentiation between items (Taylor et al., 2011). The semantics of the 627 object concepts from the property norms can be represented across 2,469 semantic features, and in the AN these correspond to the 2,469 nodes. The AN was based on Cree et al. (2006), and was trained to activate the correct pattern of semantic features from the inputs from the DNN (full details in Devereux et al., under review). The network was trained using continuous recurrent back-propagation through time over 20 processing time-ticks. As input to the AN, we took the activation over the 4,096 nodes of fc7, and reduced this to 60 dimensions using SVD (Note, RDMs calculated on the full-dimensional fc7 and the SVD-reduced layer were highly correlated (Spearman’s rho = 0.98) indicating no substantial information loss). After training, over the 20 time-ticks, the semantic features associated with the concept are gradually activated, with the speed of activation depending on the relationship to the visual features and the statistical regularities between features (i.e. whether a certain combination of features predicts the occurrence of another feature). Thus, early features to activate are shared features and visual features, followed by non-visual and distinctive features. For further implementation details see Cree et al (2006) and Devereux et al., (under review).

Like with the visual DNN, many of the 19 layers of the AN (discounting the input layer) are highly correlated and so were combined. Using k-means clustering, the 19 layers could be described well by 2 principal groups, as shown by positive silhouette values. Clustering solutions with 1,3,4 or 5 groups all contained negative silhouette values showing that 2 provided the most optimal number of clusters. After PCA, layers 1-5 were concatenated and layers 6-19 were concatenated. The dissimilarity between AN activity for different object images was calculated as 1 - Pearson’s correlation, giving an early semantic RDM and a late semantic RDM. An additional semantic RDM was created based on the concatenation of all 19 layers.

#### RDMs from time-frequency signals

Object dissimilarity from MEG signals was based on oscillatory phase patterns from source-localised data. Five regions of interest were specified covering locations known to be sensitive to visual and semantic object properties. Each ROI was specified by a coordinate and radius of 20 mm. The occipital pole (MNI: −10, −94, −16), left pVTC (MNI: −50, −52, −20), right pVTC (52, −56, −16), left ATL (MNI: −30, −6, −40) and right ATL (MNI: 30, −4, −42).Coordinates were defined based on local maxima of source localised activity to all objects. Within each ROI (defined by the center coordinate and radius), single trial activity was extracted for each vertex. Instantaneous phase was calculated for each trial and for every vertex using Morlet Wavelets using the timefreq function in EEGLAB. Phase was extracted between −700 ms and 1000 ms in 20 ms time steps, and between 4 and 95 Hz in 50 logarhythmically spaced frequency steps. A 5-cycle wavelet was used at the lowest frequency, increasing to a 15 cycle wavelet at the highest. This produced a time-frequency representation (TFR) for every trial at every vertex location in the ROI. RDMs between object TFRs were calculated at each time/frequency point, using the circular distance (Berens, 2009) between vectors of phase information over space (vertices) and over 60 ms.

For analyses at distinct frequency bands rather than at every frequency, the oscillatory RDMs were averaged across frequencies. The frequencies within each band were defined using hierarchical clustering in sensor space. TFRs were computed for each MEG sensor which were averaged across all trials and participants to produce a grand-average for each sensor. A vector was created for each frequency that included all time-points and sensors concatenated, before hierarchical clustering of frequencies using correlation as the distance measure. The resulting distances were visualised as a dendrogram to define the boundaries of the different bands. This resulted in theta (4-9 Hz), alpha (9-15 Hz), beta (16-30 Hz) and gamma (30-95 Hz).

#### RSA statistics

Each RDM based on oscillatory phase signals was correlated with the RDMs from the computational models using Spearman’s correlation. This resulted in TFRs that captured the relationship between phase information and the visual and semantic network models. RSA TFRs were calculated for each layer, ROI and participant. Random effects analysis testing for positive RSA effects was conducted for each time-frequency point using one-sample t-tests against zero (alpha 0.01). Cluster-mass permutation testing was used to assign p-values to clusters of significant tests (Maris and Oostenveld, 2007), and a maximum cluster approach was used to control for multiple comparisons across time, frequency, ROI and model RDM (Nichols and Holmes, 2002). For each permutation, the sign of the TFR correlations was randomly flipped for each participant before one-sample t-tests of the permuted data. The same permutation was applied to all ROIs and model RDMs, and the cluster with the largest mass across all analyses (sum of t-values) was retained. The p-value for each cluster in the original data was defined as the proportion of the 10,000 permutation cluster-masses (plus the observed cluster-mass) that is greater than or equal to the observed cluster-mass.

#### Peak RSA effects

To determine when different kinds of information were present relative to one another, we determined when the peak effects occurred across different regions for different visual and semantic RDMs. This analysis was performed for RSA effects within each frequency band, in addition to peak effects collapsing across frequencies. The latency of the peak was defined as the maximum Spearman’s correlation value between 50 and 500 ms. A latency was found for each frequency band, model RDM and ROI. Linear mixed effects (LME) models were used to test the relationship between the peak latency, and ROI and computational model layer. The peak frequency analysis was based on RSA effects determined across the full frequency spectrum. The peak was defined as the frequency of the maximal RSA effect between 50 and 500 ms. In both latency and frequency analyses, linear mixed effects (LME) models (using fitlme) were used to test the relationship between the peaks, and ROI and computational model layer. For visualisation, peaks are plotted as probability density functions, and using gramm (https://doi.org/10.5281/zenodo.59786).

### Granger Causality

Finally, we tested the causal relationships between RSA effects seen for visual and semantic properties and across different ROIs. Specifically, we used Granger Causality (GC) analysis to test if RSA time-courses in one region have a subsequent impact on RSA time-courses in other regions. To aid interpretability, GC analysis was applied to the RSA time-courses averaged across frequency bands and for the concatenated visual and concatenated semantic model RDMs. GC was calculated between the five ROIs and the two RSA time-courses (10 time-series in total). Each time-series was the RSA effect between 50 and 500 ms concatenated across participants. Time domain GC used the multivariate granger causality toolbox (Barnett and Seth, 2014), with a model order of 2 (40 ms) as indicated by the AIC for model order estimation. Multiple comparisons correction used FDR and an alpha of 0.05.

## Results

Our primary goal was to test how the VVP represents increasingly complex visual and semantic information over time. To achieve this, we used RSA to test if the visual and semantic information, extracted from the computational models, was represented in spatio-temporal patterns of oscillatory phase from source localised MEG signals. We analysed data from five ROIs covering occipital, posterior ventral temporal cortex and the anterior temporal lobe (ATL), that are known to be primarily implicated in the processing of visual and semantic object properties.

Time-frequency RSA (TF RSA) showed that neural patterns based on oscillatory phase had a significant relationship to both the visual and semantic models. The effects were concentrated in the first 500 ms, and seen across theta, alpha and beta frequencies (Table 1). We first present a brief overview of visual and semantic effects, before a more detailed follow-up analysis of the timing and frequencies of the effects.

**Table 1.** TF RSA results

| **Models** | **ROI** | **Freqs** | **Times** | **Mass** | **max cluster p** |
| --- | --- | --- | --- | --- | --- |
| Visual-layer 1 | Occip | 4-50 Hz | 0-730 ms | 4857 | **0.0002** |
| Visual-layer 1 | LpVTC | 4-15 Hz | 0-610 ms | 1392 | **0.0073** |
| Visual-layer 1 | RpVTC | 4-15 Hz | 0-350 ms | 1364 | **0.0077** |
| Visual-layer 1 | LATL | 4-14 Hz | 0-410 ms | 1379 | **0.0077** |
| Visual-layer 1 | RATL | 7-15 Hz | 10-370 ms | 369 | **0.0365** |
| Visual-layer 1 | RATL | 4-6 Hz | 70-350 ms | 233 | 0.1001 |
| Visual-layer 2 | Occip | 4-34 Hz | 0-750 ms | 5343 | **0.0002** |
| Visual-layer 2 | LpVTC | 4-16 Hz | 0-630 ms | 1522 | **0.0051** |
| Visual-layer 2 | RpVTC | 4-21 Hz | 0-770 ms | 2093 | **0.0035** |
| Visual-layer 2 | LATL | 4-14 Hz | 0-390 ms | 1378 | **0.0077** |
| Visual-layer 2 | RATL | 4-15 Hz | 0-750 ms | 1298 | **0.0079** |
| Visual-layer 3 | Occip | 4-32 Hz | 0-910 ms | 5599 | **0.0001** |
| Visual-layer 3 | LpVTC | 4-23 Hz | 0-690 ms | 2301 | **0.0030** |
| Visual-layer 3 | RpVTC | 4-23 Hz | 0-730 ms | 3033 | **0.0018** |
| Visual-layer 3 | LATL | 4-18 Hz | 0-590 ms | 1955 | **0.0036** |
| Visual-layer 3 | RATL | 4-17 Hz | 0-690 ms | 1943 | **0.0036** |
| Early-semantic | Occip | 5-14 Hz | 0-690 ms | 701 | **0.0145** |
| Early-semantic | LpVTC | 4-6 Hz | 170-450 ms | 251 | 0.0843 |
| Early-semantic | RpVTC | 12-21 Hz | 110-390 ms | 231 | 0.1014 |
| Early-semantic | LATL | 4-8 Hz | 0-450 ms | 584 | **0.0181** |
| Late-semantic | LpVTC | 4-8 Hz | 0-630 ms | 585 | **0.0181** |
| Late-semantic | LATL | 4-7 Hz | 0-390 ms | 447 | **0.0262** |

### Visual models

TF RSA effects were seen for all three visual models in each ROI across theta, alpha and beta frequencies (Figure 2). The strongest effects were in the occipital ROI, and were significantly greater for visual-layer 2 and visual-layer 3 models compared to visual-layer 1 (layer 2 vs. layer 1: t = 4.56, p < 0.001; layer 3 vs. layer 1: t = 3.69, p = 0.004) with no difference between visual-layer 2 and 3. In higher regions along the ventral stream, visual-layer 3 had a significantly greater fit than both layer 2 (LpVTC: t = 4.05, p = 0.002; RpVTC: t = 4.30, p = 0.001; LATL: t = 3.35, p = 0.006; RATL: t = 3.64, p = 0.004), and layer 1 (LpVTC: t = 4.90, p < 0.001; RpVTC: t = 7.00, p < 0.0001; RATL: t = 7.35, p < 0.0001) except in the LATL. These results are in line with predictions that later regions along the ventral stream represent more complex visual object information, that is in turn better captured by later layers of the visual DNN.

**Figure 2.**
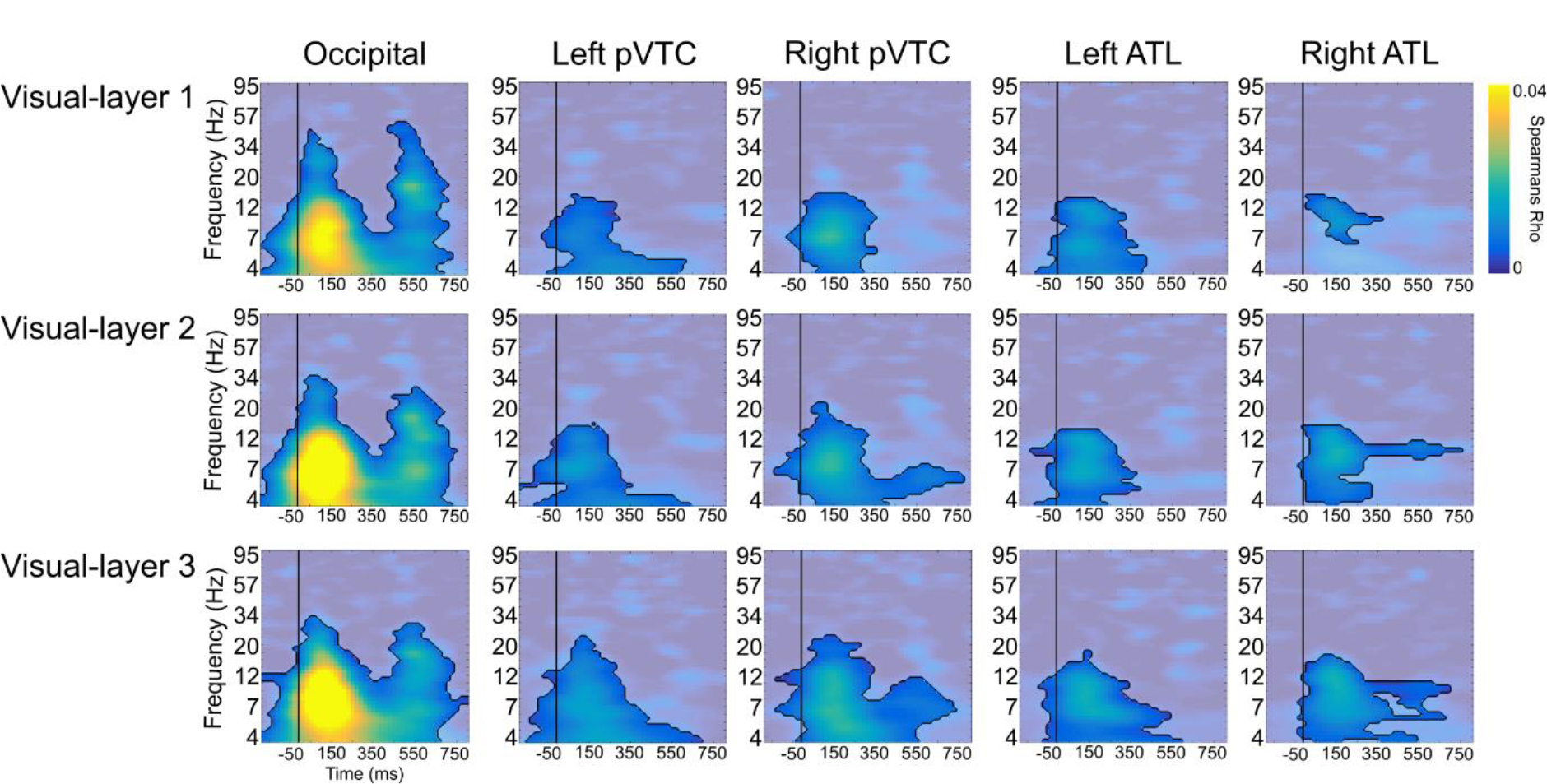
Time-frequency RSA effects of the visual DNN. Each plot shows the Spearman’s correlation values between an RDM from the visual DNN and each ROI. Significant clusters are shown outlined in black, using a threshold of p < 0.01 at the pixel level, and p < 0.05 at the cluster level (corrected for all ROIs and model RDMs tested). Non-significant time-frequency points are displayed in the background.

### Semantic models

TF RSA analysis of phase showed significant effects for both semantic models (Figure 3). Both the early-semantic and late-semantic models were significantly related to spatio-temporal phase patterns in the LATL in theta frequencies during the first 400 ms. Further, the early-semantic model had a significantly better fit compared to the late-semantic model in LATL (t = 2.96, p = 0.013). The early-semantic model was also significantly related to occipital phase patterns in theta and alpha frequencies. Finally, the late-semantic model was significantly related to the LpVTC in theta frequencies. These results show that semantic information about objects is captured through oscillatory phase patterns in the ventral stream, with the most prominent effects in theta in the pVTC and ATL – key regions supporting object semantic information over time (Clarke and Tyler, 2014; Clarke et al., 2011, 2015).

**Figure 3.**
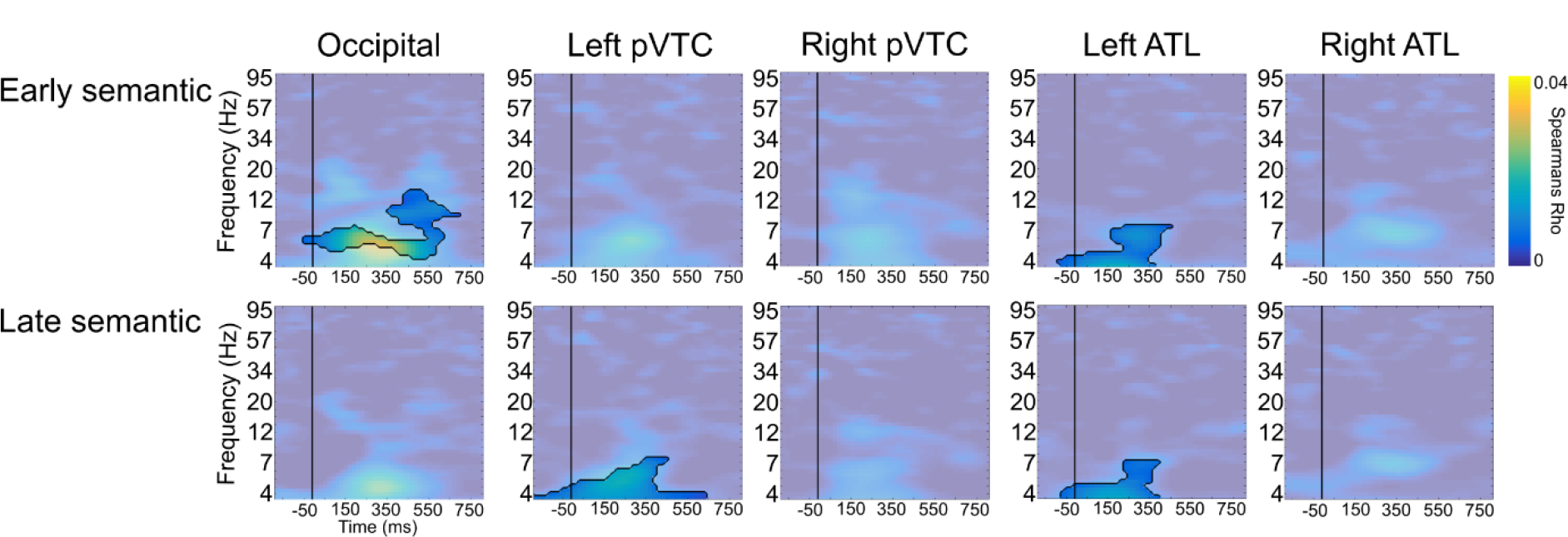
Time-frequency RSA effects of the semantic AN. Plots shows the Spearman’s correlation values between a semantic AN RDM and each ROI. Significant clusters are shown outlined in black, using a threshold of p < 0.01 at the pixel level, and p < 0.05 at the cluster level (corrected for all ROIs and model RDMs tested). Non-significant time-frequency points are displayed in the background.

### Representational changes over time

Our analysis so far shows that the combined visual DNN and semantic attractor network model are capturing neural processes along the VVP. We next sought to determine the relative changes in object information over time and region. However, it is difficult to establish when different forms of information are present based on the onsets of significant effects, due to the temporal smearing wavelet convolution creates - especially at lower frequencies - and the use of a spatio-temporal sliding window that further contributes to a smoother pattern. Further, onsets can not be easily compared across different frequencies as the temporal smearing is greater at lower compared to higher frequencies. Therefore, to determine when different kinds of information are present relative to one another, we analysed when the peak effects occurred across different regions for different visual and semantic models. We used linear mixed effects (LME) models to test the relationship between the peak time of RSA effects, and the layer of the computational model (modelled from visual layer-1, layer-2, layer-3, early-semantics, late-semantics) and hierarchical cortical level of the VVP (occipital, pVTC, ATL; where left and right hemispheres are combined).

We found a significant effect of cortical level, in that later levels of the VVP had later peak RSA effects (Beta coefficient: 26 ms (SE = 6.7 ms), t = 3.90, p = 0.0001; Figure 4A), and a significant effect of computational model layer in that later layers had significantly later peaks (Beta coefficient: 30 ms (SE = 3.6 ms), t = 8.34, p < 0.0001; Figure 4B). Further, as shown in Figure 4B, there was a prominent separation in the timing of visual and semantic peak effects. A subsequent LME model combined the data within visual and semantic models, and showed that semantic effects lagged visual effects by an estimated 88 ms (t = 8.12, p < 0.0001).

**Figure 4.**
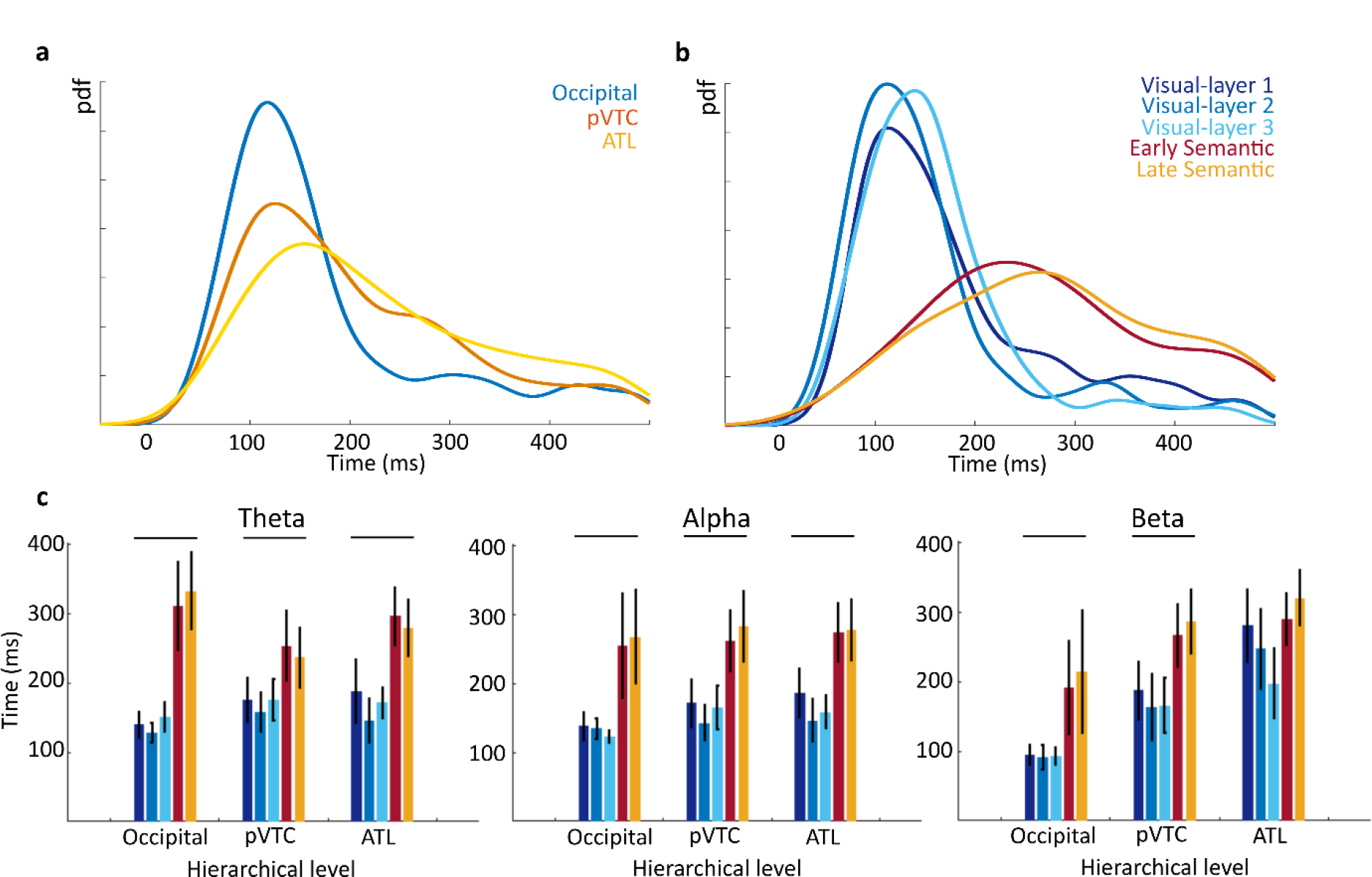
Temporal peaks for the visual DNN and semantic AN RSA effects. (A). Probability density plot showing that the latencies of the peak RSA effects follow the hierarchical levels of the VVP (data combined across hemispheres and model RDMs). (B). Probability density plot showing that the latencies of the peak RSA effects for different model RDMs have a clear distinction between visual DNN and semantic AN latencies, while later model layers tend to have later peaks. (C-E). Mean peak latencies for different model RDMs at each hierarchical level for three frequency bands where significant RSA effects were present. Plots show a general increase in latency across the models from visual to semantic (colours match those in panel b). Horizontal lines indicate a significant linear relationship between latency and model layer.

After establishing this broad pattern where effects are later in time for higher regions of the VVP and for later layers of the visual-to-semantic model, we next tested for region-specific changes in the latency of peak RSA effects within three frequency bands that showed significant effects – theta, alpha and beta. Separate LME models were run for each cortical level of the VVP for each frequency band. Significant positive effects of model layer were seen in the occipital and pVTC for theta, alpha and beta, while the ATL showed significant positive effects in theta and alpha (Figure 4C-E). This establishes that later layers of the combined computational model showed later peak RSA effects in theta, alpha and beta frequencies across all levels of the VVP, supporting our broad results of a temporal transition from visual to semantics over time in accordance with the changes seen over the successive layers of the computational model.

### Representational changes over frequency

We next tested how the peak frequency of the RSA effects changed. Using a LME model, we found a marginal effect of hierarchical cortical level (Beta coefficient = 2.2 Hz (SE = 1.1), t = 1.89, p = 0.06), but not of model layer (p = 0.21). The interaction between level and model was trending towards significance (t = 1.95, p = 0.053). To explore the interaction, separate LME models were run for each hierarchical level testing for an effect of model layer. Only the ATL showed a significant effect of model layer, where later layers of the visual-to-semantic model had lower peak frequencies (Beta coefficient: −0.74 Hz, t = 2.02, p = 0.046; Figure 5). As shown in Figure 5, plotting the probability density across frequencies suggests semantic models have median peak frequency around 5-6 Hz, while visual models have peaks closer to 10 Hz showing an alpha-theta distinction between vision and semantics.

**Figure 5.**
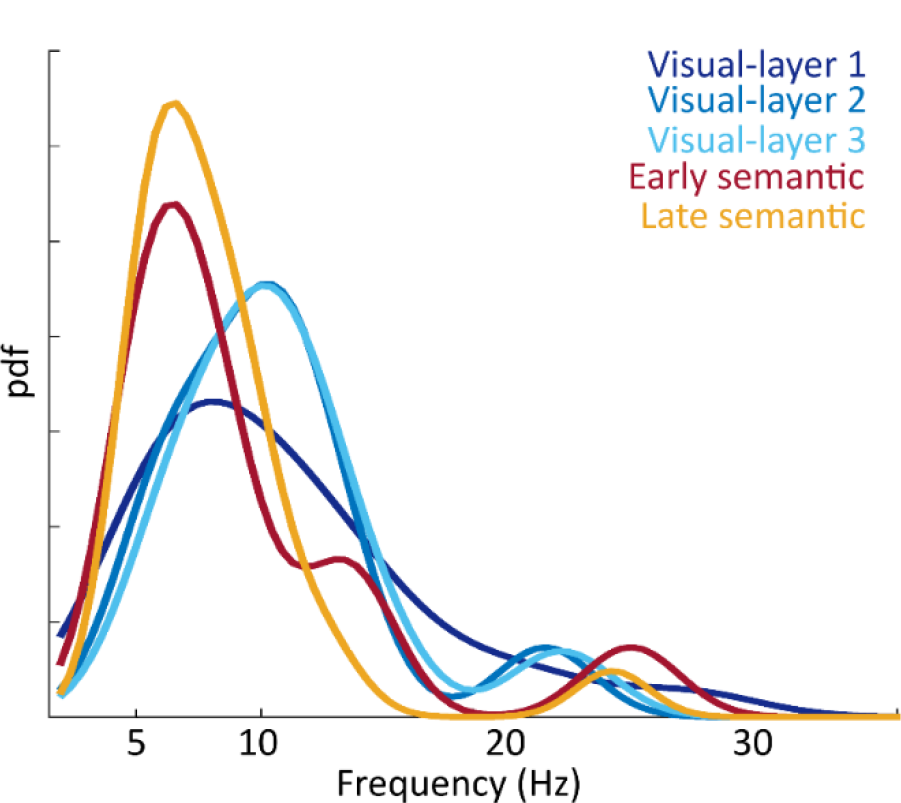
Spectral peaks for the visual DNN and semantic AN RSA effects in the anterior temporal lobe. Probability density plot showing that the peak frequency of RSA effects shows a clear distinction between visual and semantic model RDMs, where visual effects peak near 10 Hz and semantic effects peak near 5 Hz.

### Direction of information flow

The results presented so far show a visual to semantic trajectory through time and space, where effects are later in time for higher regions of the VVP, and for later layers of the visual-to-semantic model. However, focussing solely on peak effects will not fully capture the ongoing dynamics, and critically, does not tell us about the causal relationships between regions or information types. To address this, we used Granger Causality (GC) analysis to test if representations in one region have a subsequent impact on representations in other regions. For example, GC with RSA time-courses allows us to test if visual information in one region has a causal impact on subsequent visual representations in a different region, or whether visual representations have a causal impact on subsequent semantic representations. GC analysis was applied to the RSA time-courses averaged across theta and alpha bands (where effects were concentrated) to test for causal relationships between visual representations across regions, semantic representations across regions, and critically, the causal relationship between visual and semantic representations both within and across regions. For this analysis, we focus on RSA effects from the combined visual and combined semantic RDMs (Figure 6).

**Figure 6.**
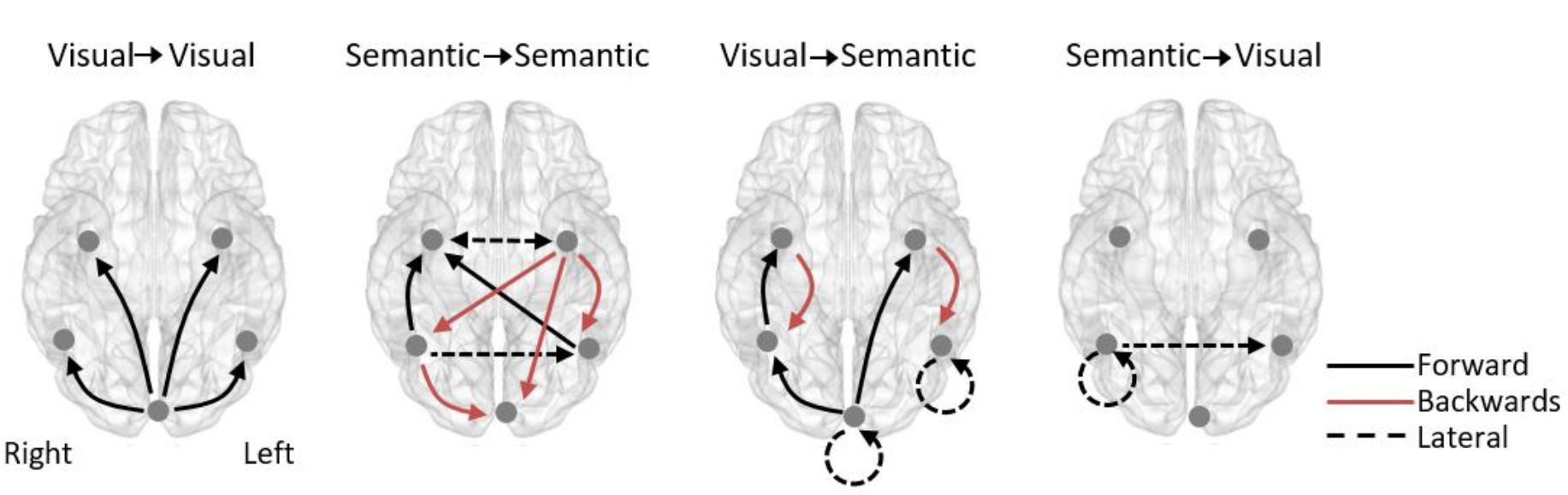
Granger causality (GC) of RSA time-courses. Images show significant GC of the RSA time-courses between regions. Each image shows how RSA effects in one region impact future RSA effects in another region. The analysis was conducted both for RSA effects across regions within the visual DNN or semantic AN model RDMs (left two images), and when the RSA effects of the visual DNN could show GC with RSA effects with the semantic AN (and vice versa; right two images). Significant connections shown using p < 0.05, FDR corrected.

We first tested how visual RSA effects impact visual effects in other regions. Significant feedforward GC was seen between the occipital region and all other regions. This suggests that visual representations in the occipital lobe have an impact on subsequent visual representations further along the the VVP in accordance with feedforward models of visual processing. Semantic RSA effects showed significant feedforward, cross-hemispheric and feedback connectivity, with both the left and right ATL playing prominent roles. Semantic effects in the LATL significantly influenced later semantic effects in more posterior regions, while the RATL showed significant connectivity from bilateral pVTC. In addition, bidirectional connectivity was seen between the ATL regions. This shows that, in contrast to visual RSA connectivity, the spread of semantic effects were associated with more complex feedforward, feedback and cross-hemispheric connectivity.

Crucially, we tested the relationships between visual and semantic RSA effects, by testing if visual RSA effects in one region influenced later semantic effects in other regions (or the same region) and vice versa. Visual RSA effects emanating from the occipital and RpVTC significantly influenced semantic effects through feedforward connectivity with the ATL, while visual RSA effects from the occipital region also influenced later semantic effects in the RpVTC. The occipital visual RSA effects influenced later semantics in the occipital, while visual effects in the LpVTC also influenced later semantic effects in the LpVTC. Finally, visual RSA effects in the ATL influenced later semantic effects in the pVTC through feedback connectivity, an effect that was present in both hemispheres. This shows a pattern where feedforward visual-to-semantic transformations occur from the occipital to LATL, and along the right VVP. Feedback visual-to-semantic transformations occurred from the ATL to pVTC bilaterally, in addition to a shifting visual-to-semantic representation within LpVTC. Lastly, semantic RSA effects had a significant effect on visual representations in the RpVTC, nd from RpVTC to LpVTC. Overall, the GC results show that feedforward processing in the VVP supports the dynamic processing of visual information, while combinations of feedforward and feedback is more central for semantics. We also highlight that visual to semantic information transitions engage feedforward and feedback connectivity, with the ATL appearing a vital region.

## Discussion

In this study, we successfully combined RSA for time-frequency phase information with a computational architecture for visual to semantic processing. Utilising a combined visual DNN and semantic AN, we were able to demonstrate how the incremental aspects of visual to semantic processes occur in the ventral stream over time, and the underlying dynamics supporting this transition. TF RSA revealed visual and semantic object properties were primarily reflected in alpha and theta activity. Spatial and temporal hierarchies were also apparent, where later layers of the computational model showed peak effects later in time, and in later regions along the posterior to anterior axis. Moreover, we also revealed more subtle dynamics underlying recognition, where feedforward connectivity supported the transfer of visual information in the VVP, and combined feedforward, feedback and intra-region dynamics supported the transition between visual and semantic information processing states. These results present the first detailed account of how oscillatory dynamics can support the emergence of meaning from visual inputs.

Here we used TF RSA with oscillatory phase information, showing that low-frequency phase carries stimulus-specific information related to visual and semantic object properties. The analysis was based on phase patterns from MEG source localised data, with our results showing that objects with more similar properties have more similar spatio-temporal phase patterns in the mass signals recorded through MEG. It is believed that the phase of low-frequency activity is suited for decoding stimulus properties for MEG, EEG and ECOG (Panzeri et al., 2015; Watrous et al., 2015a), supported by a number of studies showing that oscillatory phase carries more information about the stimulus than power (Lopour et al., 2013; Ng et al., 2013; Schyns et al., 2011). While not presented here, we also see a similar pattern with our data. While neural mass activity can be difficult to relate to the underlying neural activity, there is some suggestion that low-frequency phase of mass signals might index the timing of the underlying neural activity and its firing (Panzeri et al., 2015; Watrous et al., 2015a). As such, our effects based on spatiotemporal phase patterns may be driven by spatiotemporal activity patterns of the mass neural populations, and further suggests that cognitively relevant properties are coded in distributed neural activity patterns in space and time. However, the relative importance of a spatial or temporal activity patterns for object properties was not be determined in this study.

Previous studies in both humans and nonhuman primates have identified category-specific phase coding of objects, where different object categories have different preferred phases associated with neural activity (Turesson et al., 2012; Watrous et al., 2015b). Here, we go beyond phase dissociations between different categories, by showing that the variability in phase information relates to variability in the stimulus properties, and is the case for both visual and semantic properties. We see that phase patterns in alpha most strongly relate to visual properties from the DNN and phase patterns in theta relate to semantics. Alpha oscillations over posterior regions are linked to the sampling of the visual environment (VanRullen et al., 2014). Alpha activity is claimed to reflect a pulsed inhibition of cortical activity, where increases in alpha power result in the inhibition of a region and decreased alpha power relates to the active engagement of a region (Jensen and Mazaheri, 2010; Klimesch et al., 2007). Reductions in alpha power occur in occipital regions following visual presentation, and so our RSA effects will coincide with reductions in alpha power. Research using combined EEG and fMRI has further shown that occipital alpha power reductions correlated with increased BOLD in downstream object processing regions (Zumer et al., 2014), and so alpha activity could organise the flow of information through the VVP, as supported through our connectivity analysis (see below). However, it is also worth noting that effects of the DNN, while peaking in alpha, were seen across theta, alpha and beta frequencies.

Both alpha and theta activity are sometimes considered to have similar roles in organising neural activity (Jensen et al., 2014; Lisman and Jensen, 2013). Both alpha and theta activity are modulated by memory, but often with opposing effects (Hanslmayr et al., 2012), and our clustering of frequencies to generate the different bands revealed separate clusters for theta and alpha. Together, this suggests a functional dissociation between theta and alpha in cortex. Theta activity in the hippocampus and medial temporal lobes is tightly linked to long term memory (Fell and Axmacher, 2011; Fell et al., 2001; Halgren et al., 2015; Sederberg et al., 2003; Staresina et al., 2012). Our theta effects for semantic object properties in the pVTC and ATL are consistent with intracranial recordings in humans from anterior IT and the PRC which show a modulation of theta activity according to the semantic category of words (Halgren et al., 2015), where it is further hypothesised that ATL structures aid the encoding of attributes in coordination with theta in the hippocampus (Fell et al., 2001; Halgren et al., 2015; Staresina et al., 2012).

One implication from our study is that different primary rhythms may encode visual and semantic properties, particularly in the ATL. The concept that different frequencies code complementary aspects of a stimulus is known as multiplexing. Using EEG, Schyns and colleagues (2011) showed that posterior electrodes coded for the eyes of a face in the beta band, and the mouth in theta, showing that different features of a face are coded in different frequencies. In our study, different object features relating to vision and semantics were represented by different frequencies in the ATL - alpha and theta. Recently, the PRC within the ATL, was shown to represent both high-level visual properties and conceptual properties of objects (Martin et al., 2018). Our evidence of visual and semantic effects in the ATL may indicate that the conjoint coding of visual and conceptual properties in the PRC could be aided through a multiplexed coding scheme, that may also be useful for integrating distinct visual information within a forming semantic representation. We can speculate that given the visual environment is sampled at an alpha rate (VanRullen et al., 2014), the slower theta dynamics for semantics could be useful to integrate semantic information from the environment over multiple alpha cycles. Further ECOG investigations will be important to highlight the specific spatio-temporal-spectral signatures for vision and semantics in the ATL, and how the alpha and theta cycles are related. These studies would also offer the opportunity to test how low-frequency phase information and high frequency activity (>100 Hz) might jointly represent object information through phase-amplitude coupling (Canolty and Knight, 2010; Jensen and Mazaheri, 2010; Jensen et al., 2014). This is supported by recent work showing that high frequency activity to different object categories occurs at different phases of a low-frequency oscillation, showing how phase-amplitude coupling could relate to phase coding (Watrous et al., 2015b).

One clear step forward provided by our study, is determining how object information across different brain areas was related (also see Goddard et al., 2016 and; Ince et al., 2015 for related approaches). This is an important step, because, although our main analyses highlight parallel hierarchies of vision to semantics and posterior to anterior regions, this is likely an over simplification of the underlying activity dynamics. By combining the RSA time-courses with Granger Causality, we were able to show how information in one region changes the state of information in another region, characterising how information flows in the VVP.

As predicted by most models of visual processing, our analysis showed visual object information was associated with feedforward connectivity, in that visual representations coded in occipital alpha phase predicted future visual representations in more anterior regions in the VVP. In contrast, the flow of semantic representation effects was feedback and cross-hemispheric, similar to previous reports of feedback activity in the VVP supporting semantic processing (Campo et al., 2013; Chan et al., 2011; Clarke et al., 2011; Poch et al., 2015; Schendan and Ganis, 2012). Crucially, this analysis enabled us to test how visual representations impact future semantic representations. This analysis showed two prominent motifs. First, visual effects in the occipital region related to subsequent semantic effects in the ATL and pVTC (feedforward), and second visual effects in the ATL related to subsequent semantic effects in the pVTC (feedback). This analysis revealed more complex dynamics than suggested when only looking at peak effects, whilst also emphasising the importance of the ATL through receiving feedforward inputs, and sending top-down signals to posterior regions.

The ATL plays a central role in many theories of semantics, with differential emphasis of lateral, polar and medial aspects of the region, which may depend on stimulus modality or task (Clarke and Tyler, 2015; Damasio et al., 2004; Grabowski et al., 2001; Mehta et al., 2016; Patterson et al., 2007; Ralph, 2014). Given the spatial specificity of MEG source localisation, we did not look to test between these positions, and focus on the general role of the extended region. However, recent fMRI work using the same DNN and semantic AN approach shows that semantic effects for visual objects are represented in the PRC (Devereux et al., under review), which is consistent with a variety of other neuroimaging and neuropsychology studies showing the semantics of visual objects is dependent on the PRC (Clarke and Tyler, 2014; Kivisaari et al., 2012; Taylor et al., 2006; Tyler et al., 2013; Wright et al., 2015). Although we do not make claims about exact localisation of ATL effects from this study, our results do provide critical new evidence that can further refine these accounts. One speculative prediction we can make regarding the ATLs role, is that it initially integrates visual signals during a feedforward alpha drive whilst activating semantic object properties. The properties, represented by theta activity, then communicated through feedback activity to the pVTC (Chan et al., 2011; Clarke, 2015), with coherent activity between the posterior and anterior regions in the VVP supporting the object-specific semantics (Clarke et al., 2011) based on top-down semantic and bottom up visual signals. Theta activity may further structure alternating modes of feedforward and feedback activity (Halgren et al., 2015), with increased recurrent activity necessary under ambiguous perceptual conditions (Schendan and Ganis, 2012). Future studies utilising ECOG or depth electrodes could begin to test these predictions.

Whilst research with time-sensitive approaches converge towards a model where the initial feedforward activation activates the visual aspects of objects, before recurrent dynamics process the specific semantics (Chan et al., 2011; Clarke, 2015; Clarke and Tyler, 2015; Halgren et al., 2015; Poch et al., 2015; Schendan and Ganis, 2012), we lack an understanding of the neuro-computational principles of how vision activates meaning. Here, we tested whether oscillatory activity could represents stimulus-specific visual and semantic object properties, and showed that visual properties were most associated with alpha phase, and semantic properties were associated with theta phase information. Further, distinct modes of connectivity underpinned the flow of information, where visual information flowed in a feedforward direction, semantics in feedback, whilst the transfer between vision and semantics relied on feedforward, feedback and intra-regional flow. Our results highlight the ATL as an important region, both in representing visual and semantic information through a multiplexed code, and for the transformation of information from visual to semantic. By combining oscillations, connectivity, RSA and computational models, we show how visual signals activate meaning taking us towards a more detailed model of object recognition.

## Acknowledgements

This work was supported by a European Research Council Advanced Investigator grant under the European Community’s Horizon 2020 Research and Innovation Programme (2014-2020 ERC Grant agreement no 669820) to LKT, and from the European Research Council under the European Community’s Seventh Framework Programme (FP7/2007-2013)/ ERC Grant agreement n° 249640 to LKT.

